# Differential expression patterns and potential regulatory function of piRNAs in western honey bee larval response to *Ascosphaera apis* infestation

**DOI:** 10.1101/2022.12.27.521978

**Authors:** Minghui Sun, Qi Long, Xiaoxue Fan, Zixin Wang, Kaiyao Zhang, Siyi Wang, Xuze Gao, Sijia Guo, Qingsheng Niu, Haibing Jiang, Dafu Chen, Zhongmin Fu, Rui Guo

**Affiliations:** College of Animal Sciences (College of Bee Science), Fujian Agriculture and Forestry University, 350002, Fuzhou, Fujian, China; Apitherapy Research Institute, Fujian Agriculture and Forestry University, 350002, Fuzhou, Fujian, China; Jilin Institute of Apicultural Research, 132000, Jilin, China

**Keywords:** honey bee, *Apis mellifera*, chalkbrood, *Ascosphaera apis*, piRNA, regulation

## Abstract

piRNAs are a class of small non-coding RNAs that play an essential part in genomic defense, as well as in modulation of gene expression and diverse biological processes such as host–pathogen interaction. However, little is known about the expression pattern of the regulatory function of piRNAs in interactions between honey bees and pathogens. In this work, on the basis of our previously obtained high-quality small RNA-seq datasets from western honey bee (*Apis mellifera*) larval guts, the differential expression profile of piRNAs in *A. mellifera* larval guts after *Ascosphaera apis* infestation was analyzed, followed by structural characterization, target prediction, and regulatory network investigation. The potential roles of differentially expressed piRNAs (DEpiRNAs) in regulating host response, especially immune response, were further discussed. In this study, 504, 657, and 587 piRNAs were respectively identified in the 4-, 5-, and 6-day-old larval guts infected by *A. apis*, with 411 (53.24%) piRNAs shared. The length distribution of these piRNAs ranged from 24 nt to 33 nt and their first base had a “C” bias, similar to piRNAs discovered in other mammals and insects. Additionally, 96, 103, and 143 DEpiRNAs were detected in the 4-, 5-, and 6-day-old comparison groups; among these, piR-ame-149736, piR-ame-1066173, and piR-ame-1125190 were the most up-regulated, while piR-ame-1202932 was the most down-regulated. There were 68 DEpiRNAs shared between these three comparison groups. The targets of the DEpiRNAs in the three comparison groups were engaged in a suite of crucial functions associated with biological processes, molecular function, and cellular components, including molecular transducer activity, biological regulation, and membrane part. These targets were also relevant to diverse vital pathways such as the phosphatidylinositol signaling system, inositol phosphate metabolism, and Wnt signaling pathway. Further investigation demonstrated that targets of DEpiRNAs were involved in three energy metabolism-related pathways, seven development-associated signaling pathways, and seven immune-relevant pathways, including lysosome and endocytosis, as well as the MAPK and Jak-STAT signaling pathways. The expression trends of five randomly selected DEpiRNAs were verified using a combination of RT-PCR and RT-qPCR. Moreover, the expression levels of six genes targeted by piR-ame-945760 were detected by RT-qPCR, with the results showing that their expression trends were the same as the expression trend of piR-ame-945760, indicative of the positive correlation between piR-ame-945760 and these targets. These results suggest that *A.apis* infestation increased the overall expression level of piRNAs and altered the expression pattern of piRNAs in *A. mellifera* larval guts. DEpiRNAs potentially participate in the *A. apis* response of the host by modulating the expression of target genes associated with energy metabolism and development, as well as cellular and humoral immune response. Our findings not only offer novel insights into *A. mellifera* larva–*A. apis* interaction, but also lay the groundwork for clarifying the mechanisms underlying DEpiRNA-regulated larval response.

## Introduction

The western honey bee (*Apis mellifera*) is widely reared in China and many other countries, and provides pollination services for a substantial quantity of wild flowers and agricultural crops, thus playing crucial economic and ecological roles. However, as a eusocial insect, *A. mellifera* is susceptible to infections by various pathogens and parasites. Among these, *Ascosphaera apis* is a obligated lethal fungal pathogen that exclusively infects bee larvae and causes chalkbrood disease, which results in a dramatic reduction of colony population and productivity (Aronstein and Murray, 2009).

Non-coding RNAs (ncRNAs) are diverse and pivotal regulators in gene expression and biological processes, from cell proliferation, differentiation, apoptosis, and autophagy to growth, development, metabolism, immune defense, and host–pathogen interaction (Toden et al., 2021). Piwi-interacting RNAs (piRNAs) are a class of small ncRNAs with a length distribution ranging from 23 nt to 35 nt (Iwasaki et al., 2015; Wang et al., 2022). piRNA was first discovered in *Drosophila* and then in mice and *Caenorhabditis elegans* (Das et al., 2008; Dai et al., 2020). Thereafter, increasing numbers of piRNAs were identified in insects such as *Drosophila* (Kotov et al., 2019), *Bombyx mori* (Katsuma et al., 2021), and *Aedes aegypti* (Joosten et al., 2021). A major function of piRNAs is to specifically bind to PIWI proteins, as well as members of the Piwi, Aub, and Ago3 families, further silencing the activities of transposons (Pek et al., 2011). Furthermore, piRNAs have been demonstrated to be involved in an array of life activities such as genome rearrangement and epigenetic regulation, and to serve as novel biomarkers and therapeutic targets for disease (Khurana et al., 2010; Ashe et al., 2012). In insects, piRNAs are suggested to participate in the modulation of cognate viral infection, metabolic homeostasis, gut development, and sex determination (Kiuchi et al., 2014; Jones et al., 2016; Wang et al., 2021; Xu et al., 2022a). Moreover, accumulating evidence has shown that piRNAs are crucial regulators in responses of insects to pathogen or parasite invasion. For instance, Feng et al. (2021) found that the differential expression of piRNAs in body fat was significantly higher than in the midgut during BmNPV infection, and provided information on the interaction between DEpiRNAs and their putative targets, which may be important during BMNPV infection. However, studies on the regulatory functions of piRNAs in interactions between honey bees and pathogens or parasites have been lacking up to now. Our previous studies indicated that various types of ncRNAs such as miRNAs and circRNAs are engaged in the responses of *A. mellifea* larvae to *A. apis* invasion (Guo et al., 2019; Ye et al., 2022). In view of the involvement of piRNAs in the immune responses of insects (Feng et al., 2021), we raise the question of whether piRNAs participate in modulating the responses of *A. mellifera* larvae to *A. apis* infestation.

In our previous work, high-quality transcriptome datasets from small RNA sequencing (sRNA-seq) of *A. apis*-inoculated and uninoculated *Apis mellifera ligustica* larval guts were obtained. Here, piRNAs in the gut tissues of *A. m. ligustica* larvae inoculated with *A. apis* were characterized, followed by investigation of the differential expression pattern of piRNA (DEpiRNA) in host response to fungal infection, as well as target prediction and analysis. The potential regulatory function of differentially expressed piRNAs (DEpiRNAs) in larval responses, especially the immune response, were further resolved. To the best of our knowledge, this is the first report of piRNA-mediated bee larval response to *A. apis* infection. Our findings not only offer new insight into the interaction between *A. mellifera* larvae and *A. apis*, but also lay groundwork for clarifying the mechanism underlying piRNA-mediated host response.

## Materials and methods

### Bee larvae and fungi

*A. m. ligustica* larvae were gained from three colonies reared in the apiary of College of Animal Sciences (College of Bee Science), Fujian Agriculture and Forestry University, Fuzhou City, China. *A. apis* was previously isolated from a fresh chalkbrood mummy of *A. m. ligustica* larvae and conserved in our laboratory (Guo et al., 2018a; 2018b).

### sRNA-seq Data Source

In a previous study, *A. apis*-inoculated 4-, 5-, and 6-day old larval gut samples (Am4T, Am5T, and Am6T groups) and corresponding inoculated 4-, 5-, and 6-day old larval gut samples (Am4, Am5, Am6 groups) were prepared and subjected to RNA extraction, cDNA library construction, and deep sequencing using the Illumina MiSeqTM platform by Genedenovo Biotechnology Co., Ltd. (Guangzhou, China). Followed by strict quality control of raw data (Guo et al., 2018c; 2019a; 2019b), the high-quality raw data produced from sRNA-seq were deposited in the NCBI Sequence Read Archive (SRA) database (http://www.ncbi.nlm.nih.gov/sra/) under the BioProject number: PRJNA408312..

### Bioinformatic prediction and analysis of piRNAs

piRNAs were predicted according to our previously established procedure (Xu et al., 2022a). Briefly, (1) the clean reads were mapped to the reference genome (assembly Amel 4.5) of *A. mellifera* to obtain mapped reads; (2) clean tags were aligned to the GenBank (Benson et al., 2018) and Rfam (Griffiths et al., 2015) databases using the Blast tool to remove rRNA, scRNA, snoRNA, snRNA, and tRNA; (3) miRNAs were filtered from the remaining clean reads; (4) sRNAs with a length distribution between 24 nt and 33 nt were screened on the basis of the length characteristics of piRNAs, and only those aligned to a unique position were retained as candidate piRNAs.

The expression level of each piRNA was normalized with the TPM method. Next, the length distribution and first base bias of piRNAs were investigated based on the prediction result. The UpSet plot and ridgeline plots of the piRNA expression levels were visualized using the OmicShare platform (https://www.omicshare.com/tools/, (accessed on 5 December 2022 and 11 December 2022)).

### Identification of DEpiRNAs

Based on the criteria of |log_2_ FC| ≥1 and *P* ≤ 0.05, DEpiRNAs in Am4 vs Am4T, Am5 vs Am5T, and Am6 vs Am6T comparison groups were screened using Edger software (Robinson et al., 2010). Venn analysis of DEpiRNAs in each comparison groups was conducted using the related tool in the OmicShare platform (https://www.omicshare.com/tools/ (accessed on 27 October 2022)).

### Prediction and investigation of DEpiRNA-targeted genes

According to the method described by Xu et al. (2022a), target prediction of DEpiRNAs was performed with TargetFinder software (Allen et al., 2005). Then, the targets were respectively annotated in the GO (https://www.geneontology.org(accessed on 27 October 2022)) and KEGG (https://www.genome.jp/kegg/(accessed on 27 October 2022)) databases to gain corresponding functional and pathway annotations. Further, on basis of the predicted targeting relationships, regulatory networks between DEpiRNAs and target genes were constructed, followed by visualization using Cytoscape v.3.3.0 software (Smoot et al., 2011).

### RT-PCR validation of DEpiRNAs

Total RNA from the gut tissues of 4-, 5-, and 6-day-old larvae were isolated with the FastPure Cell/Tissue Total RNA Isolation Kit v2 (Vazyme), followed by evaluation of purity and concentration using a Nanodrop 2000 spectrophotometer (Thermo Fisher, Waltham, MA, USA). One DEpiRNA (piR-ame-945760) in the Am4 vs. Am4T comparison group, two (piR-ame-1186994, piR-ame-904316) in the Am5 vs. Am5T comparison group, and two (piR-ame-978292, piR-ame-1199278) in the Am6 vs. Am6T comparison group were randomly selected for RT-PCR validation. Specific stem-loop primers, as well as upstream primers (F) and universal downstream primers (R), were designed by using DNAMAN software and then synthesized by Sangon Biotech Co., Ltd. (Shanghai, China). Using HiScript 1st Strand cDNA Synthesis Kit (Yeasen), reverse transcription was performed with stem-loop primers and the obtained cDNAs were used as templates for PCR amplification, which was conducted on a T100 thermocycler (Bio-Rad, Hercules, CA, USA) under the following conditions: pre-denaturation at 95 °C for 5 min, 40 amplification cycles of denaturation at 95 °C for 10 s, annealing at 55 °C for 30 s, and elongation at 72 °C for 1 min, followed by a final elongation step at 72 °C for 10 min. The PCR system (20 μL) included 1 μL of diluted cDNA, 10 μL of PCR mix (Yeasen), 1 μL of forward primers, 1 μL of reverse primers, and 7 μL of DEPC water. The amplification products were checked on 1.8% agarose gel electrophoresis with Genecolor (Gene-Bio, Shenzhen, China) staining.

### RT-qPCR verification of DEpiRNAs and target genes

The above-mentioned five randomly selected DEpiRNAs were also subjected to RT-qPCR detection, which was conducted on an Applied Biosystems QuantStudio 3 system (Thermo Fisher, Waltham, MA, USA) with the conditions: pre-denaturation step at 95 °C for 5 min, 40 amplification cycles of denaturation at 95 °C for 10 s, annealing at 60 °C for 30 s, and elongation at 72 °C for 15 s, followed by a final elongation step at 72 °C for 10 min. The reaction system included 1.3 μL of cDNA, 1 μL of forward primers, 1 μL of reverse primers, 6.7 μL of DEPC water, and 10 μL of SYBR Green Dye. All reactions were performed in triplicate. The snRNA *U6* gene (GenBank ID:725641) was selected as the inner reference. The relative expression level of each DEpiRNA was calculated following the 2^-ΔΔCt^ method (Livak and Schmittgen, 2001).

Based on the target prediction and annotation results, six genes (sorting nexin-6 (*SNX6*, GeneID: 727452); suppressor of cytokine signaling 5 (*SOCS5*, GeneID: 552036); V-type proton ATPase subunit d (*ATP6V1D*, GeneID: 409946); E3 ubiquitin-protein ligase TRIM37-like (*E3UBL-TRIM37*, GeneID: 413376); ubiquitin-conjugating enzyme E2 Q2 (*UBE E2 Q2*, GeneID: 409607); AP-1 complex subunit gamma-1 (*AP1G1;* GeneID: 412884)) associated with five immune pathways—endocytosis, the Jak-STAT signaling pathway, oxidative phosphorylation, ubiquitin-mediated proteolysis, and lysosome—were selected for RT-qPCR determination. Specific forward primers (F) and reverse primers (R) were designed with DNAMAN software and then synthesized by Sangon Biotech Co., Ltd. (Shanghai, China). According to the instructions of the SYBR Green Dye kit (Vazyme), the reactions were performed on an ABI QuantStudio 3 fluorescence quantitative PCR instrument with the conditions: pre-denaturation at 95 °C for 5 min, denaturation at 95 °C for 10 s, and annealing and extension at 60 °C for 30 s for 40 cycles. The reaction system contained 10 μL of SYBR Green Dye,1.3 μL of cDNA template, 1 μL of forward and reverse primers, and 6.7 μL of DEPC water. The *actin* gene (GenBank ID:36320842) was selected as the inner reference. All reactions were performed in triplicate. The relative expression level of each DEpiRNA was calculated using the 2^-ΔΔCt^ method. Detailed information about the primers used in this work was presented in Table S1.

### Statistical Analysis

Statistical analyses were conducted with SPSS software (IBM, Amunque, NY, USA) and GraphPad Prism 7.0 software (GraphPad, San Diego, CA, USA). Data are shown as mean *±* standard deviation (SD) and were subjected to Student’s *t*-test. Fisher’s exact test was performed using R software 3.3.1 to screen significant (*p* < 0.05) GO terms and KEGG pathways.

## Results

### Number, characteristics, and overall expression level of piRNAs in *A. m. ligustica* larval guts inoculated with *A. apis*

Here, 504, 657, and 587 piRNAs were discovered in the Am4T, Am5T, and Am6T groups, respectively **(Figure 1)**. After removing redundant ones, a total of 772 *A. m. ligustica* piRNAs were identified. Additionally, 411 piRNAs were shared by the three groups, while the numbers of unique ones were 27, 66, and 114, respectively **(Figure 1)**.

**FIGURE 1.**
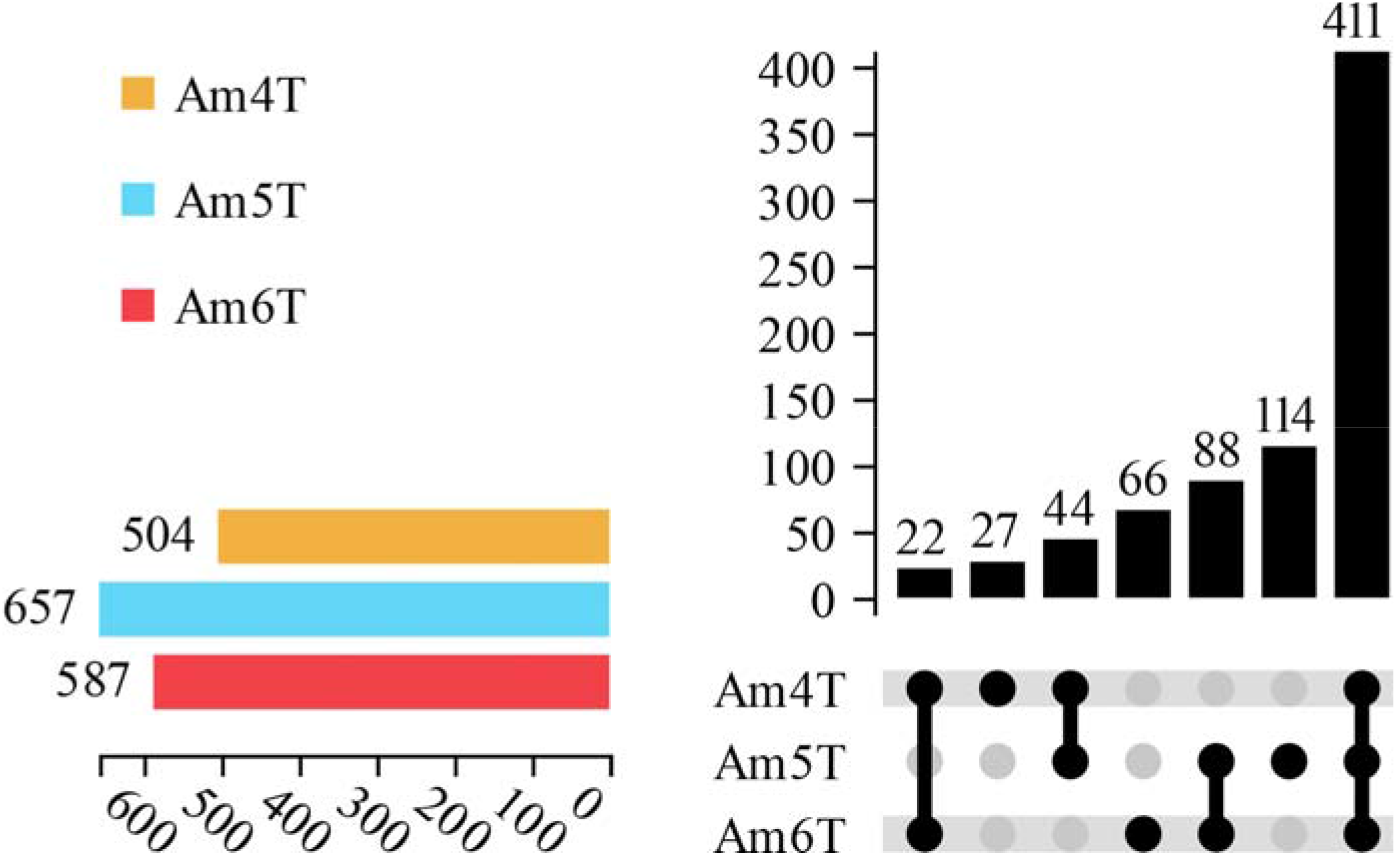
Differential expression of piRNAs in *A. m. ligustica* larval guts inoculated with *A. apis*. The lower-left color columns indicate the numbers of piRNAs in different groups; the lower-right nodes indicate the piRNAs shared by the three groups; the connections among different groups indicate the shared piRNAs, while the teamless nodes indicate the unique piRNA in each group.

Additionally, the length distribution of the identified piRNAs in *A. apis*-inoculated larval guts ranged from 24 nt to 33 nt (**Figure 2A, C, E**). Furthermore, the first base of these piRNAs had a “C” bias (**Figure 2B, D, F**).

**FIGURE 2.**
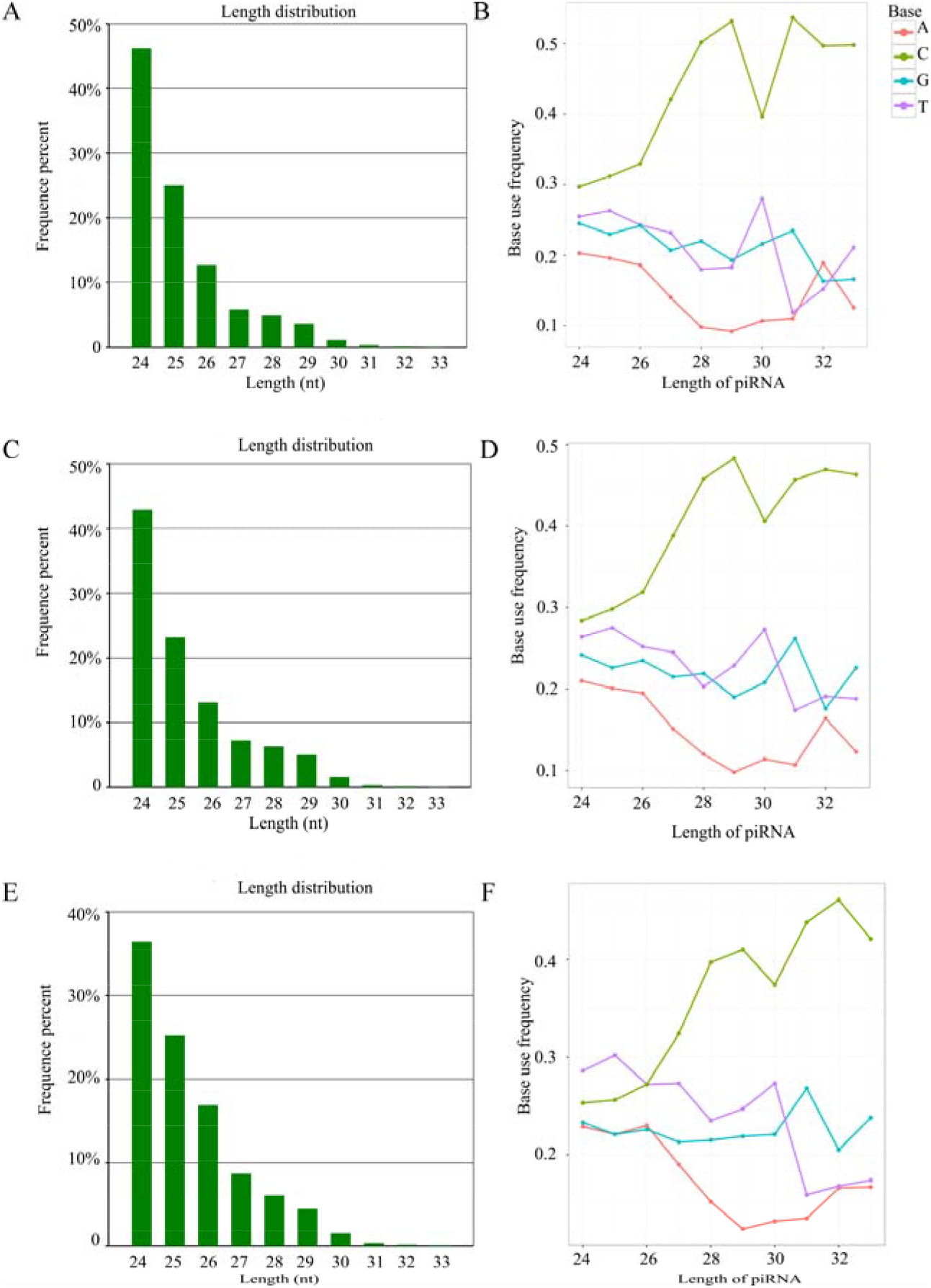
Length distribution and first base bias of the identified piRNAs in *A. apis*-inoculated larval guts. **(A-B)** Length distribution and first base bias of piRNAs in Am4T group; **(C-D)** Length distribution and first base bias of piRNAs in Am5T group; **(E-F)** Length distribution and first base bias of piRNAs in Am6T group.

### Differential expression pattern of DEpiRNAs in *A. m. ligustica* larval guts after *A. apis* infestation

Further investigation suggested that the overall expression levels of piRNAs among three *A.apis*-inoculated or uninoculated groups were similar, whereas the overall expression level of piRNAs in the *A. apis*-inoculated larval gut was higher than in the uninoculated larval gut (**Figure 4A**). In the Am4 vs. Am4T comparison group, eight up-regulated piRNAs and 88 down-regulated ones were identified, of which the most up-regulated piRNA was piR-ame-149736 (log_2_ FC = 12.30, *P* = 2.24 × 10^-7^), followed by piR-ame-220719 (log_2_ FC = 1.13, *P* = 0.024), and piR-ame-403890 (log_2_ FC = 1.13, *P* = 0.024), while the three most down-regulated piRNAs were piR-ame-1202932 (log_2_ FC = −21.37, *P* = 3.21 × 10^-8^), piR-ame-1188062 (log_2_ FC = −19.97, *P* = 1.10 × 10^-8^), and piR-ame-1188284 (log_2_ FC = −18.19, *P* = 3.51 × 10^-7^). Comparatively, 26 up-regulated piRNAs and 77 down-regulated ones were detected in the Am5 vs. Am5T comparison group; among these, the most up-regulated piRNA was piR-ame-1066173 (log_2_ FC = 2.09, *P* = 0.017), followed by piR-ame-506267 (log2 FC=1.96, *P*=0.018) and piR-ame-788843 (log2 FC=1.96, *P*=0.018), whereas the three most significantly down-regulated piRNAs were piR-ame-1202932 (log2 FC = −22.19, *P* = 6.65 × 10^-8^), piR-ame-1188062 (log2 FC = −20.33, *P* = 1.44 × 10^-7^), and piR-ame-1184482 (log_2_ FC = −19.21, *P* = 3.20 × 10^-7^). In the Am6 vs. Am6T comparison group, 61 up-regulated piRNAs and 82 down-regulated ones were discovered; of these, the three most up-regulated piRNAs were piR-ame-1125190 (log_2_ FC = 13.32*,P* = 7.01 × 10^-5^), piR-ame-982406 (log_2_ FC = 12.32, *P* = 0.00013), and piR-ame-810373 (log_2_ FC = 12.30, *P* = 1.12 × 10^-5^), while the most down-regulated piRNA was piR-ame-1202932 (log_2_ FC = −22.05, *P* = 1.07 × 10^-8^), followed by piR-ame-1188062 (log_2_ FC = −20.26, *P* = 4.71 × 10^-11^) and piR-ame-1184482 (log_2_ FC = −19.29, *P* = 4.04 × 10^-10^). Moreover, Venn analysis showed that a total of 68 piRNAs were shared by these three comparison groups; the numbers of unique ones were 22, 20, and 56, respectively (**Figure 4E**).

**FIGURE 4.**
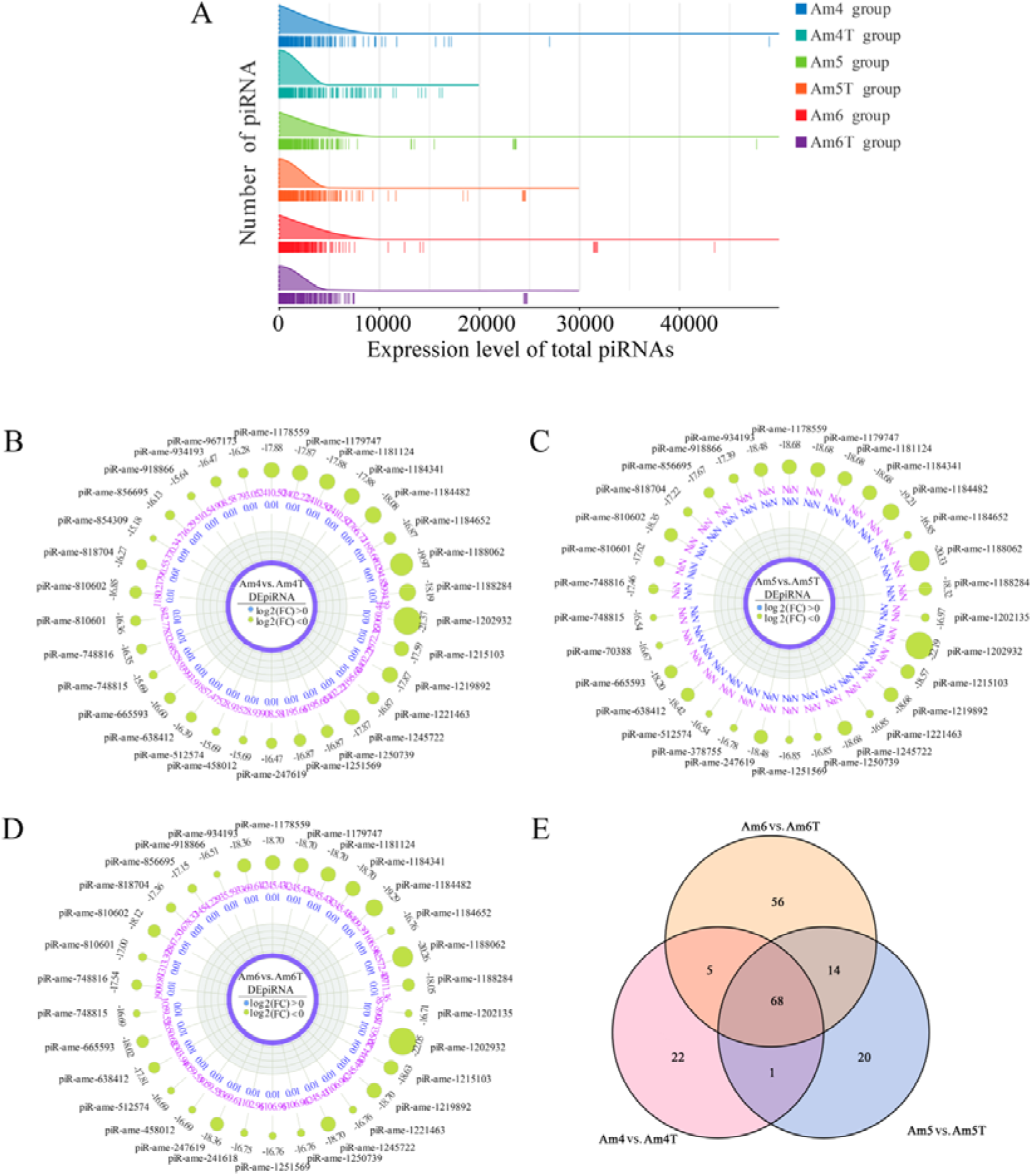
Ridgeline plots of overall expression levels of piRNAs in *A.apis*-inoculated and uninoculated *A. m. ligustica* larval guts **(A)**. Radar maps of the top 30 in the Am4 vs. Am4T, Am5 vs. Am5T, and Am6 vs. Am6T comparison groups **(B-D)**. Venn diagram of DEpiRNAs in the three comparison groups mentioned above **(E)**.The peak of the ridge indicates the most abundant piRNA in that group, and each vertical line indicates a piRNA.

### Annotation and analysis of DEpiRNA-targeted genes

DEpiRNAs in the 4-day-old comparison group was predicted to target 11517 genes, which can be annotated to 48 GO terms, including 20 biological process-related terms such as biological regulation and cellular processes, 11 molecular function-related terms such as molecular transducer activity and signal transducer activity, and 17 cellular component-related terms such as extracellular matrix and synapse part (**Figure 5A**). In contrast, the DEpiRNAs in the 5-day-old comparison group can target 11255 genes, which were involved in 20 biological process-associated terms such as biological regulation and signaling, 11 molecular function-associated terms such as molecular transducer activity and signal transducer activity, and 19 cellular component-associated terms such as extracellular matrix and synapse part (**Figure 5B**). In addition, DEpiRNAs in the 6-day-old comparison group could target 12077 genes, which were engaged in 20, 11, and 17 terms related to biological process, molecular function, and cellular component, respectively (**Figure 5C**).

**FIGURE 5.**
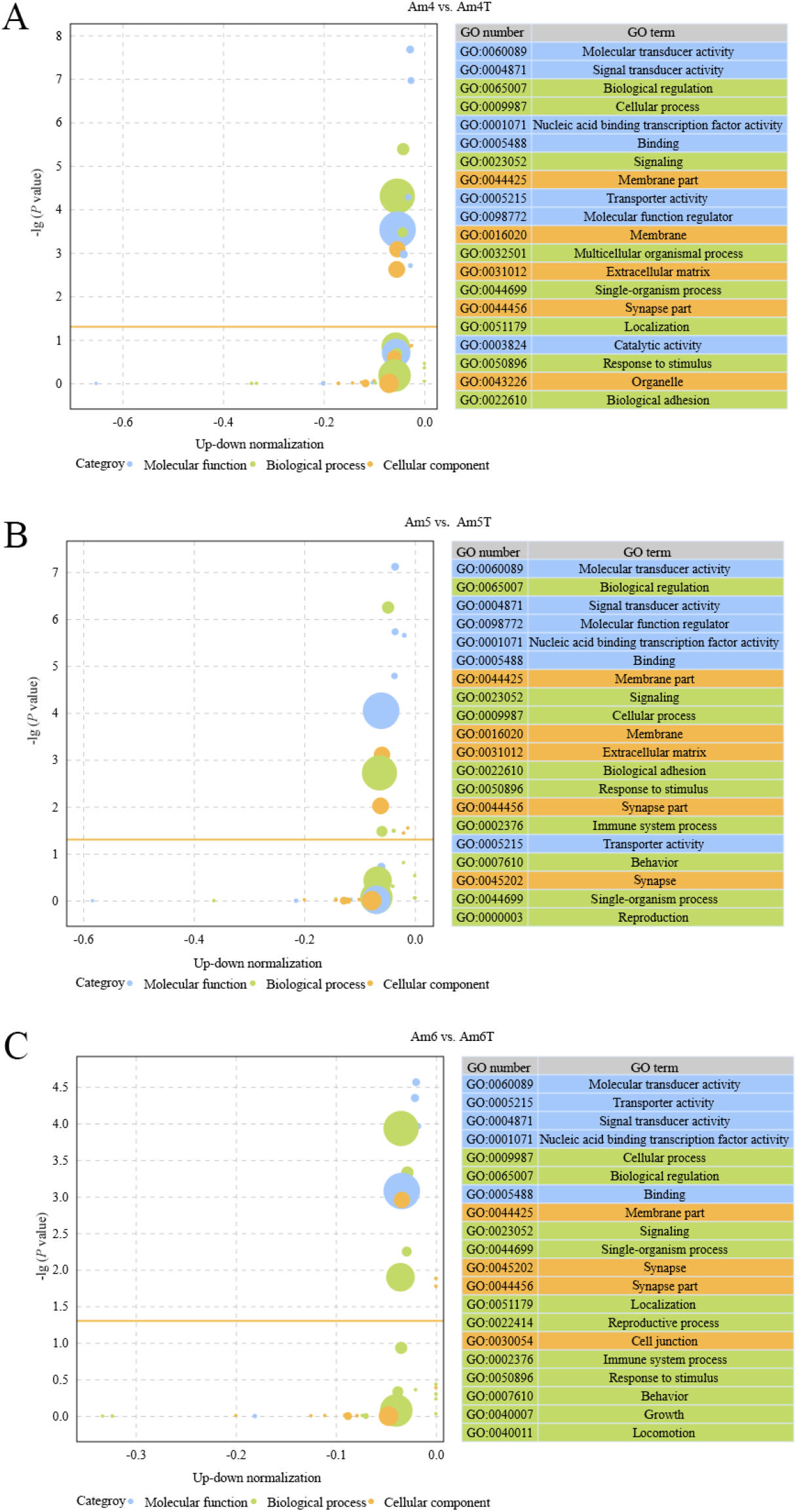
Bubble diagrams of top 20 GO terms annotated by target genes of DEpiRNAs in the Am4 vs. Am4T **(A)**, Am5 vs. Am5T **(B)**, and Am6 vs. Am6T **(C)** comparison groups. Circle size represents the number of targets; the larger the circle size, the greater the number of targets.

DEpiRNAs in the 4-day-old comparison group were annotated to 151 KEGG pathways including lysosome, the Notch signaling pathway, and oxidative phosphorylation **(Figure 6A)**; Comparatively, DEpiRNAs in the 5-day-old comparison group were involved in 145 pathways including endocytosis, the Hippo signaling pathway, and nitrogen metabolism **(Figure 6B)**. Additionally, DEpiRNAs in the 6-day-old comparison group were relevant to 151 KEGG pathways including the Jak-STAT signaling pathway, the Wnt signaling pathway, and sulfur metabolism **(Figure 6C)**.

**FIGURE 6.**
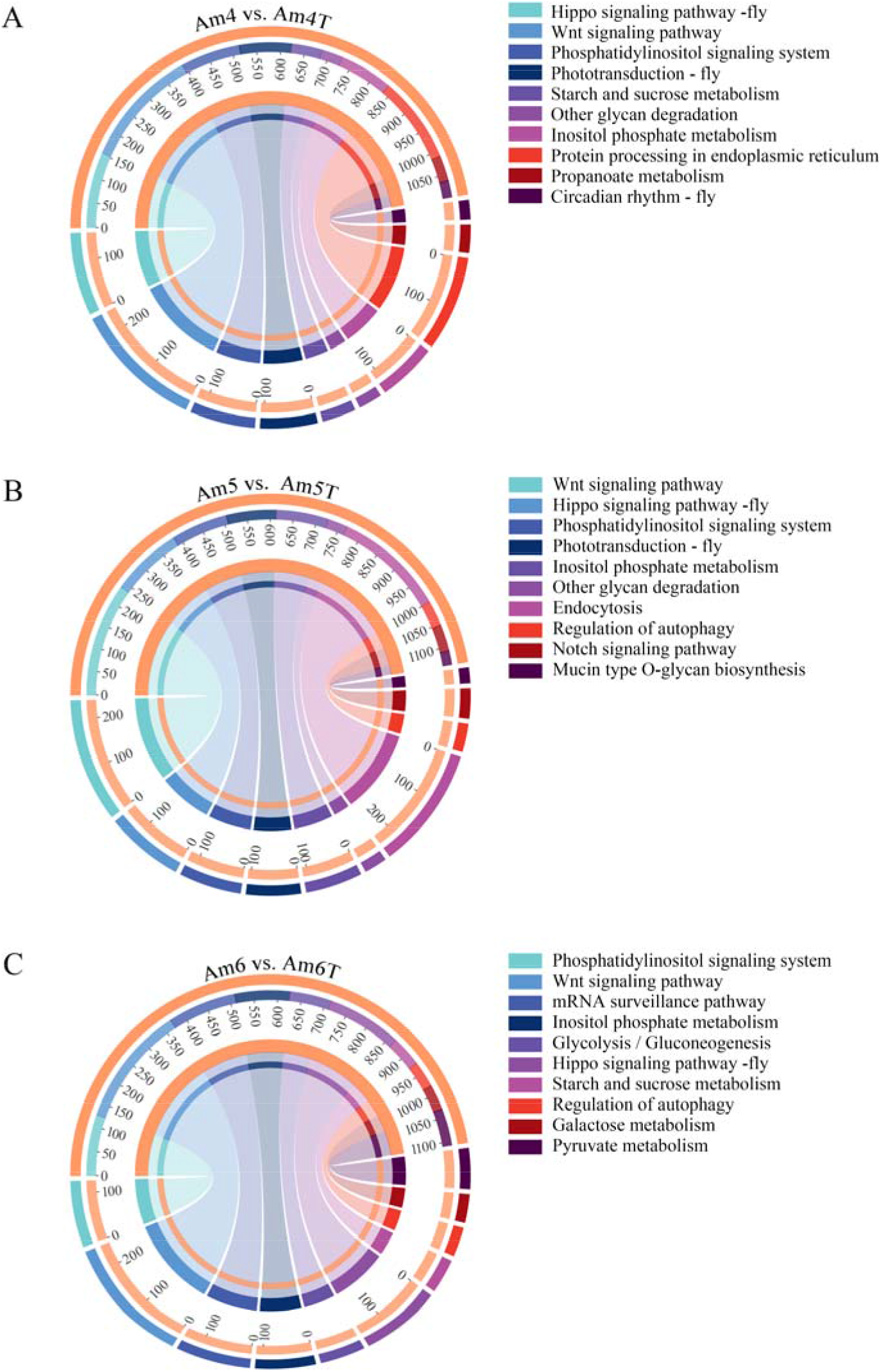
String diagrams of KEGG pathways annotated by target genes of DEpiRNAs in the Am4 vs. Am4T **(A)**, Am5 vs. Am5T **(B)**, and Am6 vs. Am6T **(C)** comparison groups.

### Regulatory networks between DEpiRNAs and targets

It was observed that complex regulatory networks formed between DEpiRNAs and corresponding targets in the above-mentioned three comparison groups, and each DEpiRNA in the aforementioned three comparison groups had more than two targets (**Figure 7, see also Table S2**). Additionally, piR-ame-785504 and piR-ame-220719 in the 4-day-old comparison group were bound to the greatest number of targets (3020 and 2779) (**Figure 7A**), piR-ame-904316 and piR-ame-608008 in the 5-day-old comparison group targeted the most genes (1919 and 1373) (**Figure 7B**), while piR-ame-854132 and piR-ame-806084 in the 6-day-old comparison group were found to link to the highest number of targets (2983 and 2700) (**Figure 7C**).

**FIGURE 7.**
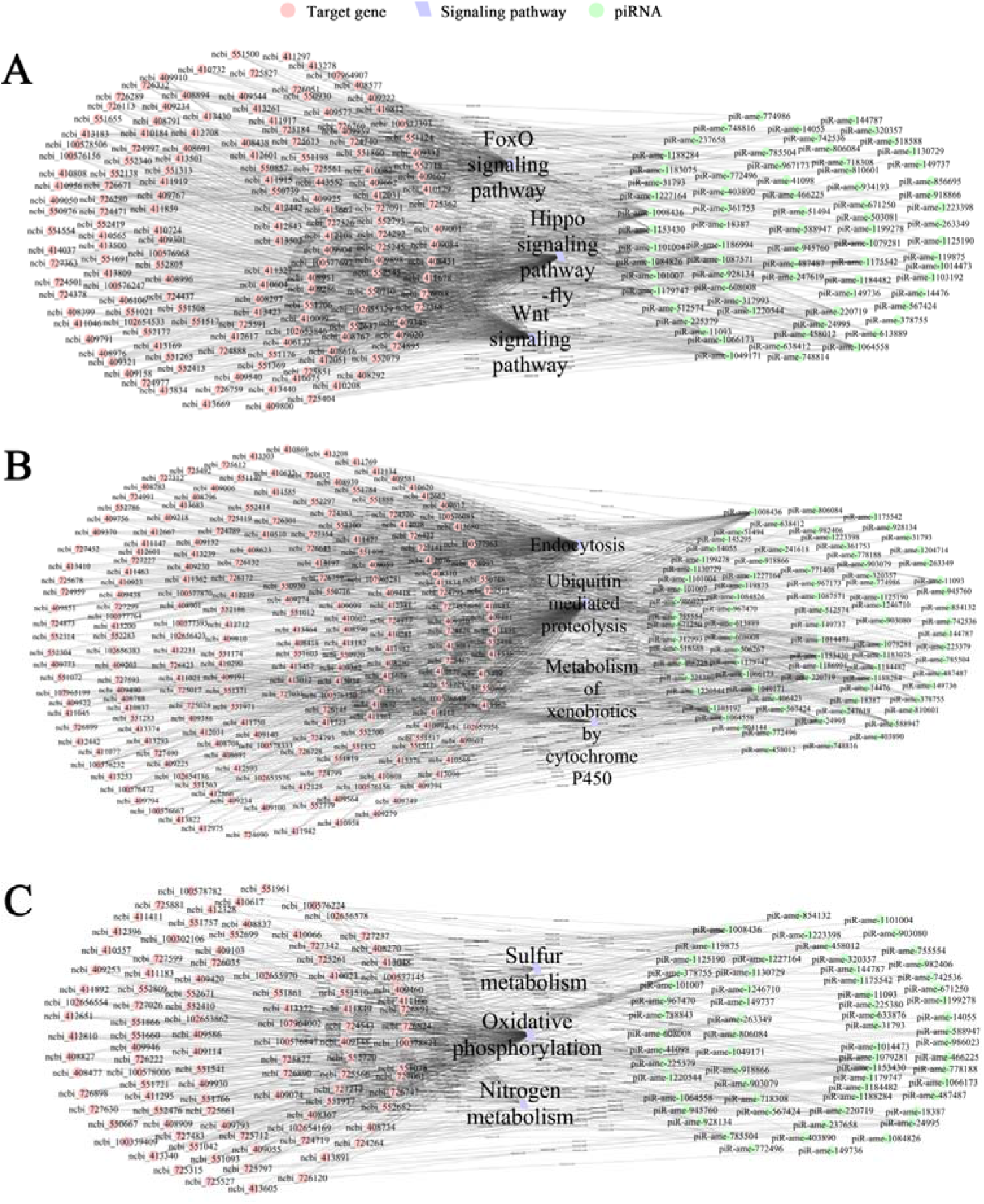
Regulatory networks between DEpiRNAs and target genes in the 4-, 5-, and 6-day-old comparison groups. **(A)** Regulatory network associated with seven development-related signaling pathways. **(B)** Regulatory network associated with immune-related pathways. **(C)** Regulatory network associated with three energy metabolism-related pathways.

It is suggested that the 226, 226, and 229 targets in these three comparison groups were engaged in seven development-associated signaling pathways: Hippo, Wnt, FoxO, Notch, mTOR, TGF-beta, the hedgehog signaling pathway, and dorso-ventral axis formation (**Figure 7A**). In addition, 300, 261, and 269 targets were involved in seven immune-relevant pathways, namely, ubiquitin-mediated proteolysis, lysosome, MAPK signaling pathway-fly, metabolism of xenobiotics by cytochrome P450, drug metabolism-cytochrome P450, endocytosis, and the Jak-STAT signaling pathway (**Figure 7B**). Moreover, 68, 61, and 85 targets were detected to be associated with three energy metabolism pathways: sulfur metabolism, nitrogen metabolism, and oxidative phosphorylation (**Figure 7C, see also Table S2**).

### RT-PCR and RT-qPCR confirmation of DEpiRNA

RT-PCR results demonstrated that fragments with the expected size (about 70 bp) could be amplified from five randomly selected DEpiRNAs, confirming the expression of these DEpiRNAs in *A. m. ligustica* larval gut.

In addition, RT-qPCR results showed that the expression trends of these five DEpiRNAs were in accordance with those in transcriptome data, confirming the reliability of the sRNA-seq datasets used in this work (**Figure 9**).

**FIGURE 8.**
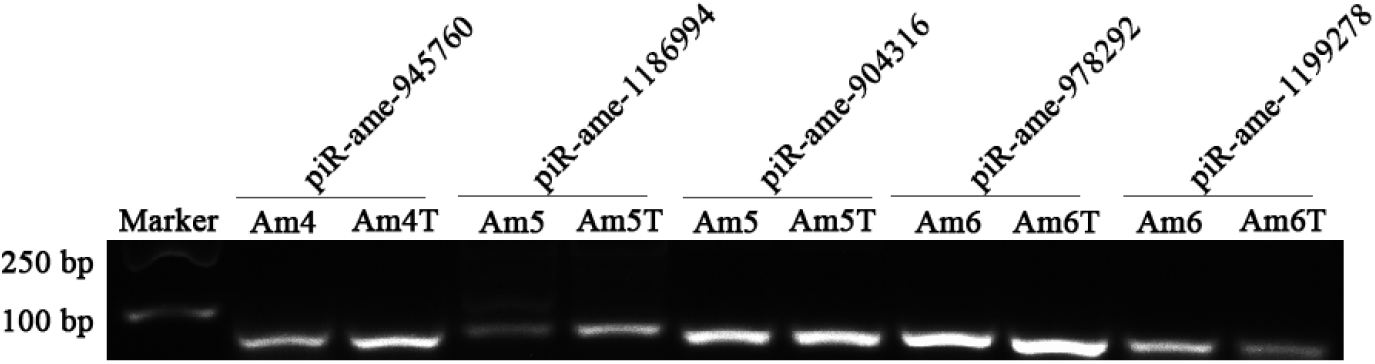
Agarose gel electrophoresis for amplification of products from five DEpiRNAs.

**FIGURE 9.**
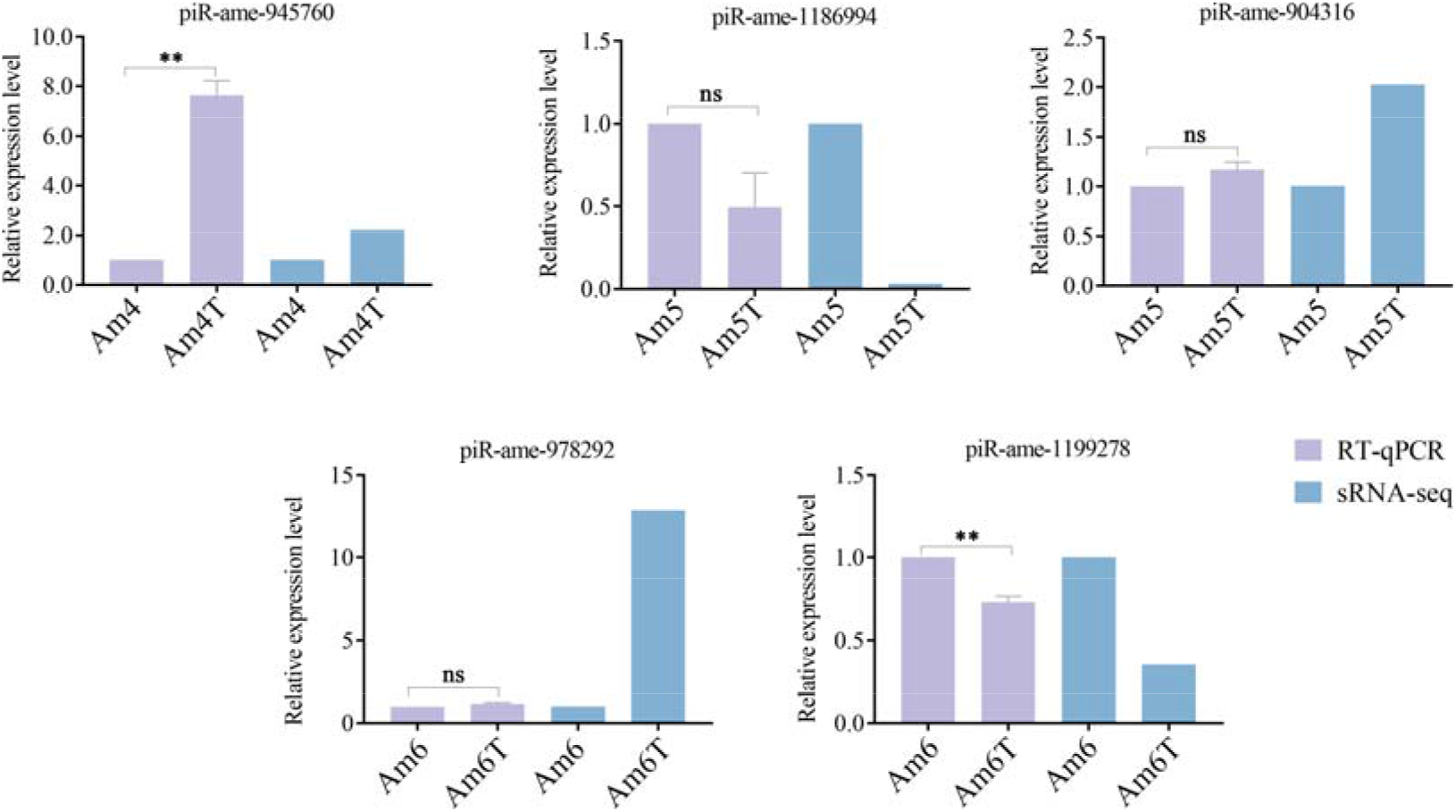
RT-qPCR detection of five DEpiRNAs. piR-ame-945760 was a DEpiRNA selected from the Am4 vs. Am4T comparison group, piR-ame-1186994 and piR-ame-904316 were two DEpiRNAs selected from the Am5 vs. Am5T comparison group, and piR-ame-978292 and piR-ame-1199278 were two DEpiRNAs selected from the Am6 vs. Am6T comparison group.** indicates *p* < 0.01.

### RT-qPCR confirmation of target genes

Six genes associated with immune-related pathways were randomly selected for RT-qPCR examination. The results demonstrated that the expression levels of these six targets were up-regulated in the *A. apis*-inoculated groups compared to the uninoculated groups, and their expression trends were same as the corresponding DEpiRNA (**Figure 10**).

**FIGURE 10.**
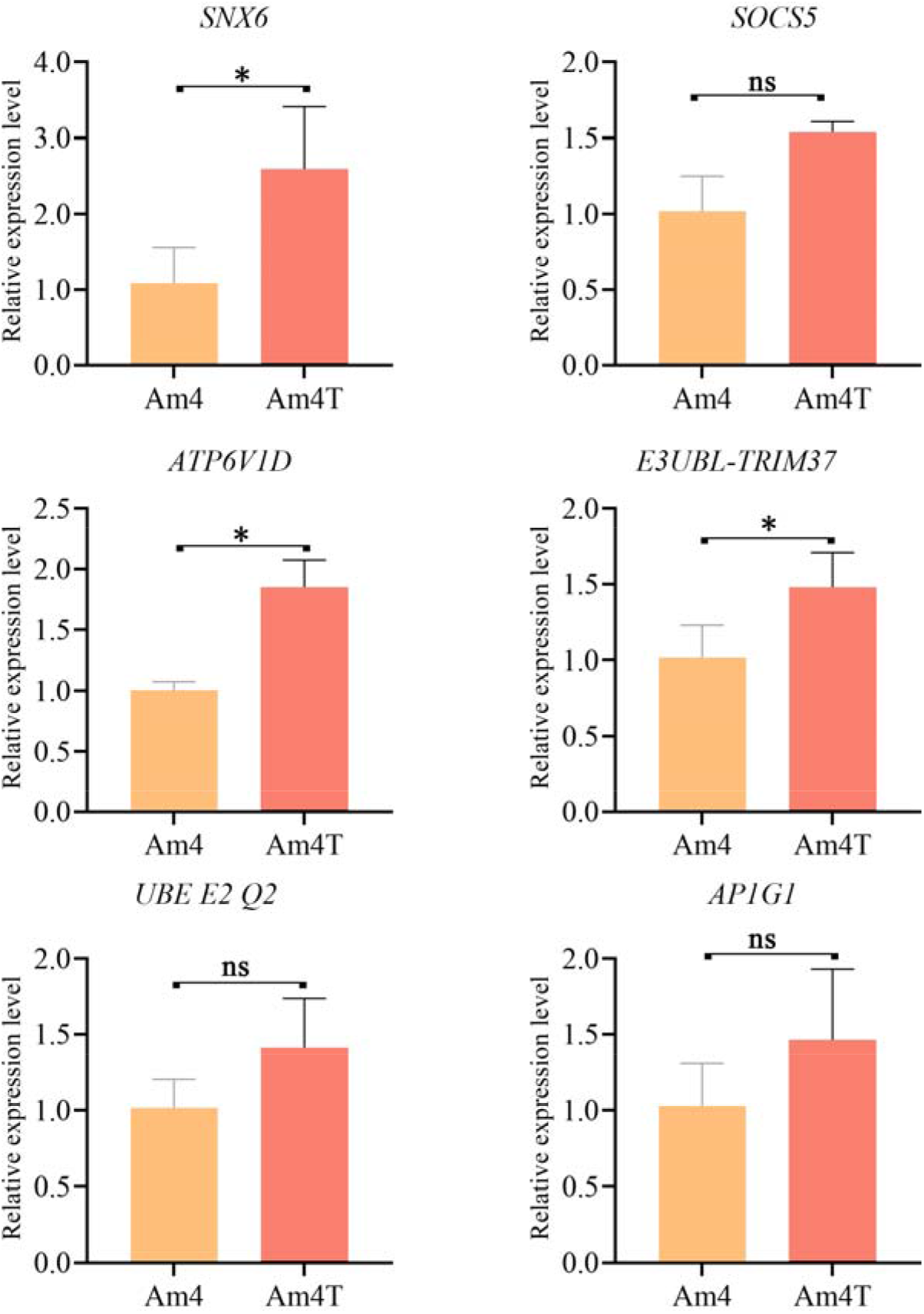
RT-qPCR determination of six target genes of piR-ame-945760. These genes were associated with endocytosis **(A)**, the Jak-STAT signaling pathway **(B)**, oxidative phosphorylation **(C)**; ubiquitin-mediated proteolysis **(D-E)**, and lysosome **(F)**. ns indicates non-significant; * indicates *p* < 0.05.

## Discussion

Though piRNAs have been reported to be in involved in interactions between mammals (Yu et al., 2019; Li et al., 2021) or insects (Wang et al., 2018) and pathogens, little was known about piRNA-involved responses of honey bees to pathogen infections. Here, based on the obtained high-quality sRNA-seq datasets, 504, 657 and 587 piRNAs were discovered for the first time in the gut tissues of *A. m. ligustica* 4-, 5-, and 6-day-old larvae challenged by *A. apis*. In addition, 411 piRNAs were shared by Am4T, Am5T, and Am6T groups, indicated that these shared piRNAs played vital roles during the *A. apis* infection process. We previously identified 775, 831, and 765 piRNAs in the uninoculated 4-, 5-, and 6-day-old larval guts (Xu et al., 2022b). It was found that 592 (62.45%) piRNAs were shared by the *A. apis*-inoculated and uninoculated larval guts, indicative of the fundamental roles of these shared piRNAs in both the development and the fungal response of *A. m. ligustica* larval guts. Combining the piRNAs identified in the *A. apis*-inoculated and uninoculated larval guts, a total of 948 non-redundant piRNAs were found. Given that the quantity of known piRNAs in honey bees at present is very limited, these piRNAs could enrich reservoir of *A. mellifera* piRNAs. Furthermore, we detected that the identified piRNAs in this work ranged from 24 nt to 33 nt in length and had a “C” bias, similar to the properties of the piRNAs identified in uninoculated *A. m. ligustica* larval guts and some other mammals such as mice (Vourekas et al., 2015). This suggests that the characteristics of *A. m. ligustica* piRNAs were unchanged under *A. apis* infection, reflecting the structural stability of piRNAs, as reported in other species (He et al., 2022; Li et al., 2022).

Increasing evidence has shown that piRNAs are involved in the responses of insects to pathogen infections. For example, Morazzani et al. (2012) discovered that arbovirus replication in *Mosquito soma* can trigger the host piRNA pathway. The silencing of piRNA-associated proteins reduced virus-specific piRNA-like molecules and enhanced viral replication and production. In the present study, we found that the piRNA pathway also has an anti-RNA virus function in lepidopteran cultured cells, as in silkworm (Katsuma et al., 2018). Previously, we found that the expression pattern of piRNAs in *A. m. ligustica* workers’ midguts was altered due to infection by *Nosema ceranae*, another prevalent bee fungal parasite (Xu et al., 2022a). In this current work, we observed that the overall expression level of piRNAs in the *A.apis*-inoculated larval guts was higher than in the uninoculated larval guts, indicating the impact of *A. apis* infection on piRNA expression at an integral level. Additionally, 96, 103, and 143 piRNAs were differentially expressed in the *A. m. ligustica* 4-, 5-, and 6-day-old larval guts after *A. apis* infection, further suggesting that the *A. apis* infection changed the expression profile of piRNAs in the larval guts. Moreover, the number of DEpiRNAs grew with infection time, indicating that more piRNAs were employed by the host in response to the fungal invasion. We can infer that this may be a strategy of the host to respond to the *A. apis* infestation. Intriguingly, a portion of piRNAs were observed to be highly up- or down-regulated during the host response, such as piR-ame-149736 and piR-ame-1202932 in the 4-day-old larval gut, piR-ame-1066173 and piR-ame-1202932 in the 5-day-old larval gut, and piR-ame-1125190 and piR-ame-1202932 in the 6-day-old larval gut infected by *A. apis*. Considering the complicated interactions between honey bee larvae and *A. apis*, these DEpiRNAs were speculated to exert crucial functions on the larval response to *A. apis* infestation, and thus deserve additional investigation in the near future, e.g., overexpression and knockdown by feeding of corresponding mimics and inhibitors (Guo et al, 2017).

In recent years, studies have demonstrated that piRNAs are pivotal regulators in gene expression and diverse life activities, in a similar way to miRNA. Gou et al. reported that the mechanism by which piRNAs act is similar to miRNAs in somatic cells, where they induce spermatogenesis through the elimination of a large number of mRNAs in mouse elongating spermatids (Gou et al., 2014). piRNA-19128 and *KIT* gene expression were observed to be negatively correlated during spermatogenesis, suggesting that piRNA-19128 plays an important regulatory role in gonadal development (Guo et al., 2018). Through overexpression and knockdown of piRNA-3312, Guo et al (2017) found that piRNA-3312 targets the gut esterase 1 gene to negatively regulate resistance to the insecticide deltamethrin. In the present study, DEpiRNAs in the 4-, 5-, and 6-day-old comparison groups were predicted to target 11517, 11255, and 12077 genes, respectively. These targets are engaged in a series of critical functions such as molecular transducer activity, biological regulation, and membranes, as well as pathways such as the phosphatidylinositol signaling system, inositol phosphate metabolism, and the Wnt signaling pathway. This is suggestive of the extensive regulatory role of DEpiRNAs in larval guts, which is in accordance with associated documentation of piRNAs in other animals, including insects (Waiho et al., 2020; Feng et al., 2021). DEpiRNAs were connected to seven immune-associated signaling pathways, including endocytosis, metabolism of xenobiotics by cytochrome P450, and ubiquitin-mediated proteolysis. In addition, the result of regulatory network analysis showed that some DEpiRNAs could target multiple immune-related genes (**Figure 7**), Altogether, our results revealed the potential regulation ability of these DEpiRNAs in host response through target immune genes.

Insects fight against pathogenic microorganisms through innate immunity (Zhang et al., 2021). Endocytosis and phagocytosis are two major cellular immune pathways in the honey bee (Aronstein and Murray, 2021). The ubiquitin-proteasome system plays a vital part in stress response, host adaptation, and fungal pathogenesis (Cao et al., 2021). Here, we observed that 83, 87, and 123 DEpiRNAs in 4-, 5, and 6-day-old *A. apis*-inoculated larval guts potentially targeted 109, 107, and 108 genes relative to endocytosis; 78, 89, and 122 DEpiRNAs targeted 83, 80, and 86 targets associated with ubiquitin-mediated proteolysis; 79, 84, and 111 DEpiRNAs targeted 63, 62, and 64 targets associated with lysosome; while 31, 41, and 48 DEpiRNAs targeted 16, 14, and 18 targets associated with metabolism of xenobiotics by cytochrome P450. The results showed that corresponding DEpiRNAs likely participate in the host cellular immune response to *A. apis* infestation by regulating the expression of target genes. Multiple immune signaling pathways, including Toll, IMD, JAK/STAT, JNK, and insulin, are engaged in insect immunity (Hillyer et al., 2016). The JAK/STAT signaling pathway is a universally expressed intracellular signal transduction pathway involved in many important biological processes, including cell proliferation, differentiation, apoptosis, and immune regulation, and has been shown to be involved in antimicrobial, antiviral, and antimalarial responses, it is the main signaling mechanism of many cytokines and growth factors (Hillyer et al., 2016; Xin et al., 2020). MAP kinase is one of the oldest and most evolutionarily conserved signaling pathways, and is critical for many immune processes, including innate immunity, adaptive immunity, and the initiation of immune responses to activation-induced cell death (Dong et al., 2002). In this work, 83, 87, and 122 DEpiRNAs in the 4-, 5-, and 6-day-old comparison groups were detected to target 70, 69, and 69 genes, which were engaged in humoral immune pathways such as MAPK and the Jak-STAT signaling pathway, suggesting the involvement of corresponding DEpiRNAs in larval humoral immune response (**Figure 7**). Together, these results reveal that DEpiRNAs are putative modulators in both cellular and humoral immune responses of *A. m. ligustica* larval guts to *A. apis* infestation.

## Conclusions

A total of 772 piRNAs were identified in the gut tissues of *A. m. ligustica* larvae infected by *A. apis*, with length distribution and first base bias similar to those of piRNAs in mammals. The *A.apis* infestation increased the overall expression level of piRNAs and changed the expression pattern of piRNAs in *A. m. ligustica* larval guts. DEpiRNAs potentially participate in the *A. apis-response* of the host by modulating the expression of target genes associated with a suite of vital pathways such as energy metabolism, development, and cellular and humoral immune response.

## Authors contribution statement

RG, ZF, and DC conceived and planned the experiments. MS, QL, XF, KZ and SW carried out the experiments and analyzed and interpreted the data. XG, SG, QN, and HJ designed the figures. QL, MS and RG reviewed and edited the paper. All authors contributed to the review and approval of the manuscript for publication.

## Acknowledgments

All authors thanks reviewers and editors for their constructive comments and recommendations. RG appreciates the love from his beloved wife and daughter.

## Supplementary Material

The Supplementary Material for this article can be found online at: https://www.frontiersin.org/ Supplementary Table 1| Detailed information about the primers of DEpiRNAs and target genes Supplementary Table 2| Detailed information about the targeting relationship between DEpiRNAs and target genes

